# Lineage Tracing Reveals Clone-Specific Responses to Doxorubicin in Triple-Negative Breast Cancer

**DOI:** 10.1101/2025.03.18.643980

**Authors:** Daylin Morgan, Andrea L. Gardner, Amy Brock

## Abstract

Triple-negative breast cancer, characterized by aggressive growth and high intratumor heterogeneity, presents a significant clinical challenge. Here, we use a lineage-tracing system, ClonMapper, which couples heritable clonal identifying tags with single-cell RNA-sequencing (scRNA-seq), to better elucidate the response to doxorubicin in a model of TNBC. We demonstrate that, while there is a dose-dependent reduction in overall clonal diversity, there is no pre-existing resistance signature among surviving clones. Separately, we found the existence of two transcriptomically distinct clonal subpopulations that remain through the course of treatment. Among clones persisting across multiple samples we identified divergent phenotypes, suggesting a response to treament independent of clonal identity. Finally, a subset of clones harbor novel changes in expression following treatment. The clone and sample specific responses to treatment identified herein highlight the need for better personalized treatment strategies to overcome tumor heterogeneity.

## Introduction

Breast cancer remains one of the most prevalent malignancies worldwide, with triplenegative breast cancer (TNBC) accounting for 15-20% of all diagnoses [1]. TNBC is defined by a lack of estrogen receptor (ER), progesterone receptor (PR), and human epidermal growth factor receptor 2 (HER2) overexpression. This subtype is notable for its aggressive nature, increased propensity for metastasis, and poorer overall clinical outcomes [2, 3]. While advancements in targeted therapies have led to significant improvements in patient outcomes for receptor-positive breast cancer subtypes, chemotherapy remains the standard of care for TNBC [4, 5].

The development of effective treatments for TNBC is significantly hindered by the substantial intratumoral heterogeneity observed within these tumors [6, 7]. A better understanding of how this heterogeneity confers increased chemoresistance at a clonal level is needed to design more efficacious treatments. Past attempts to characterize underlying clonal responses to chemotherapy in TNBC have been limited to allelic frequency measurements of bulk populations and molecular subtyping [8, 9]. However, recent advancements in lineage-tracing technologies [10–12] and single-cell RNA sequencing (scRNA-seq) [13–17] now enable the tracking and characterization of clonal responses to treatment with high granularity. Combining these techniques provides a powerful approach to understanding how tumor cell populations may adapt to and evade treatment.

To investigate the clone-specific response of TNBC to treatment with doxorubicin, we utilized ClonMapper, a DNA barcode lineage-tracing platform [10, 18]. ClonMapper introduces stably integrated expressed nucleotide barcodes, enabling the longitudinal quantification of clonal identity within a population. Importantly, this feature enables the pairing of clone-specific lineage information with detailed transcriptomic profiles, allowing for the comprehensive analysis of clonal evolution during selection and recovery following chemotherapy exposure. This study reveals significant clonal selection in the absence of a transcriptomically defined pre-existing resistance signature in a TNBC model cell line. Acute treatment with doxorubicin triggered distinct, clone- and sample-specific cell state changes, highlighting the critical need for personalized treatment strategies that account for the unique clonal composition and characteristics of individual patient tumors.

## Results

### Doxorubicin reduces clonal diversity in barcoded MDA-MB-231 cells

A barcoded MDA-MB-231 cell line was established using the ClonMapper lineage tracing system [18] (Fig. 1a). MDA-MB-231 cells were transduced at a low multiplicity of infection (0.1) with a ClonMapper viral barcode library of unique 20-nucleotide DNA barcodes and tagBFP reporter. Following transduction, 1000 BFP^+^ cells were isolated by fluorescence-activated cell sorting (FACS) and expanded to ~5 × 10^6^ cells before sequencing to determine barcode diversity (426 unique barcodes; Fig. 1b). A single archive population (hereafter referred to as Pretreatment) was expanded to ~20 × 10^6^ and used for all subsequent studies. The initial barcoded MDA-MB-231 cell library, was ~90% BFP+ (Fig. S1a). Barcoded MDA-MB-231 cells were split into 18 replicate pools of 5 × 10^4^ cells. Replicate pools were treated with doxorubicin (250, 400, and 550 nM, n=6) for 48 hours before drug removal and washout. Following treatment, there was a marked decrease in viability across all doses, with one sample each of 400nM and 550 nM failing to recover from treatment. Treated populations were maintained in culture until the cell population reached approximately 5 × 10^6^ cells. Following recovery from treatment, genomic DNA was harvested from each recovered sample to assess clonal diversity. Illumina sequencing of integrated DNA barcodes revealed a dose-dependent loss of diversity (Fig. 1b,c; Fig. S1c,d). Expression of BFP in each post-treatment samples was measured by flow cytometry to normalize total barcoded cell abundances (Fig. S1a). One sample (550-5) had no remaining expression of BFP and likely no barcoded clones remaining following treatment. This lack of barcodes in post-treatment samples may be attributed to high survivorship of the rare non-barcoded clones present in the naïve population (~10%).

**Fig. 1.**
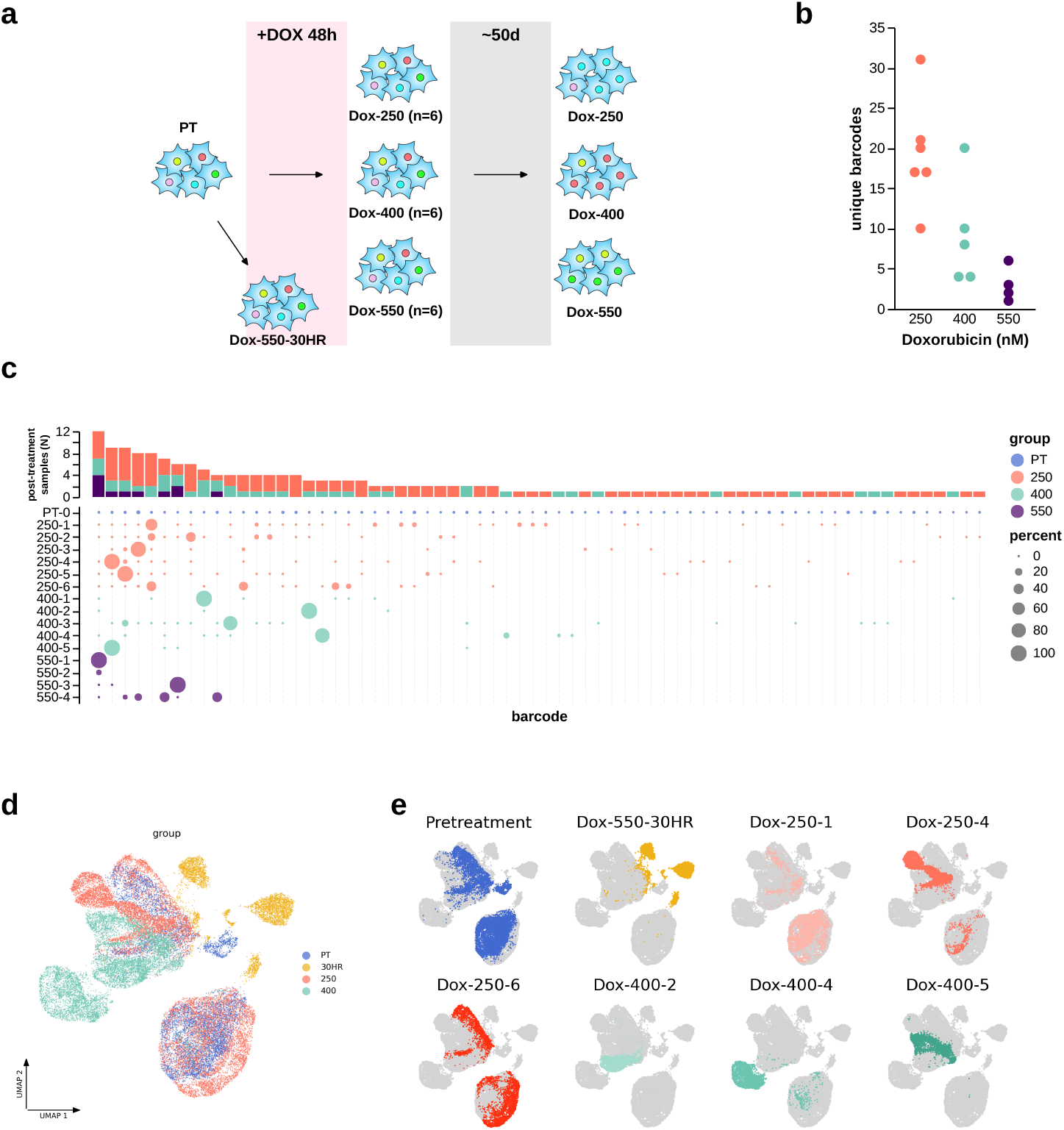
Barcoded MDA-MB-231 clonal diversity decreases with increasing dose of doxorubicin. **a)** Experimental workflow for selection and sampling of a barcoded TNBC cell line, for scRNA-seq and targeted barcode-seq. **b)** Total unique barcodes following parallel treatment of barcoded MDA-MB-231 cells at 250nM (n=6), 400 nM (n=5), 550 nM (n=4). **c)** Dot plot of barcode abundance, dots are scaled by percent abundance within a single population, barcodes are ordered by total number of samples it is detected in then cumulative abundance (bottom). Total number of post-treatment samples where barcode is detected (top). **d)** UMAP from scRNA-seq of barcoded MDA-MB-231 cells. Annotated by sample group, Pretreatment (n=1), Dox-550-30HR (n=1), Dox-250 (n=3), Dox-400 (n=3) **e)** UMAP from scRNA-seq of Barcoded MDA-MB-231 colored by each sample respectively.

In 11/15 recovered samples, greater than 90% of the barcoded cell population was comprised of less than three unique clones per sample. 31 clones are found in more than one sample, of these 7 are highly abundant (>90%) in 1-2 samples (Fig. S1b). Given this selection of unique clones within a population of MDA-MB-231 cells, we next wanted to determine whether transcriptomic state prior to treatment biased clonal survival. Using the ClonMapper barcoding system we performed clonally-resolved single-cell RNA-sequencing of 8 cell samples: the cell library prior to treatment (PT), three samples each that were treated with 250 nM and 400 nM doxorubicin respectively (250-1, 250-2, 250-6; 400-2, 400-4, 400-5), and a split from the naïve population that was exposed to 550 nM doxorubicin for 30 hours (Dox-550-30HR) (Fig. 1a). We analyzed 33,115 total cells with median unique molecular identifier (UMI) of 73,226 and median genes per cell of 8,954 (Fig. 1d,e). To resolve clonal identity of individual cells, we extracted expressed ClonMapper barcodes for each cell from unmapped reads. These reads were first length corrected to account for 3’ bias in sequencing by comparing sequences <19bp with all sequences 19-21 bp in length (Fig S3a). Following this, barcodes were corrected for sequencing error using message-passing clustering with starcode [19] and filtered on known cell library sequences (Fig S3b). Barcode reads were confidently assigned to 88.59% of the cells with each population having consistently lower total identified barcodes (Fig S3c). Lower total unique barcodes compared to amplified genomic sequences is expected given the sampling limitations of scRNA-seq.

### Barcoded MDA-MB-231s harbor pre-existing and distinct clonal subpopulations

To determine the overall relationship between transcriptomic and clonal identity we performed Leiden clustering [20] (Fig. S4a) and hierarchical clustering (Fig. S4b) on the basis of clonal identity. Cross-tabulating these classifications revealed two preexisting clone-specific subpopulations within the barcoded MDA-MB-231 cell line (Fig. 2a,b). As has been previously noted [21], these transcriptomic states are defined by the presence or absence of ESAM expression (Fig. 2f). We found a strong correlation between transcriptomic identity of these subpopulations across treatment and samples (Fig. S4c). Top differentially expressed genes characterizing these subpopulations within their Pretreatment clone-clusters were consistently expressed across samples (Fig. 2e). Within the 6 post-treatment samples, 20 unique clones were detected and four of these were identified in at least two samples (Fig. S3c). The relatively low throughput (<10000 cells per sample) of single-cell RNA-sequencing limited our ability to profile lowly-abundant clones, particularly from post-treatment samples. The stability of ESAM expression in distinguishing the underlying clones of these subpopulations was confirmed through successive rounds of isolation via FACS followed by barcode sequencing (Fig. 2g). These results support prior evidence suggesting that ESAM separates highly distinct subpopulations in MDA-MB-231 cells with no interconversion [21]. When these cells were maintained in standard culture conditions alongside recovering populations there was a steady drop-off in overall clonal diversity and a marked decrease in the presence of clone-cluster 1 (Fig. 2c,d; Fig S2a). The total number of passages a given clone remained detectable within the population was positively associated with initial measured abundance (Fig. S2b). Assuming a population level doubling time of 27 hours and exponential growth we computed approximate growth rates of all clones in normal conditions. Clone-cluster 2 had higher growth rates overall compared to clone-cluster 1 (Fig S2c). Clonal abundance, particularly of samples in doses 400 and 550 nM were representative of relatively higher growth rate clones (Fig S2d). If clones have stable growth rates we would expect that faster growing clones alongside regular down-sampling would slowly bias the population. It is possible that modified culture conditions would preferentially select clone-cluster 1, instead.

**Fig. 2.**
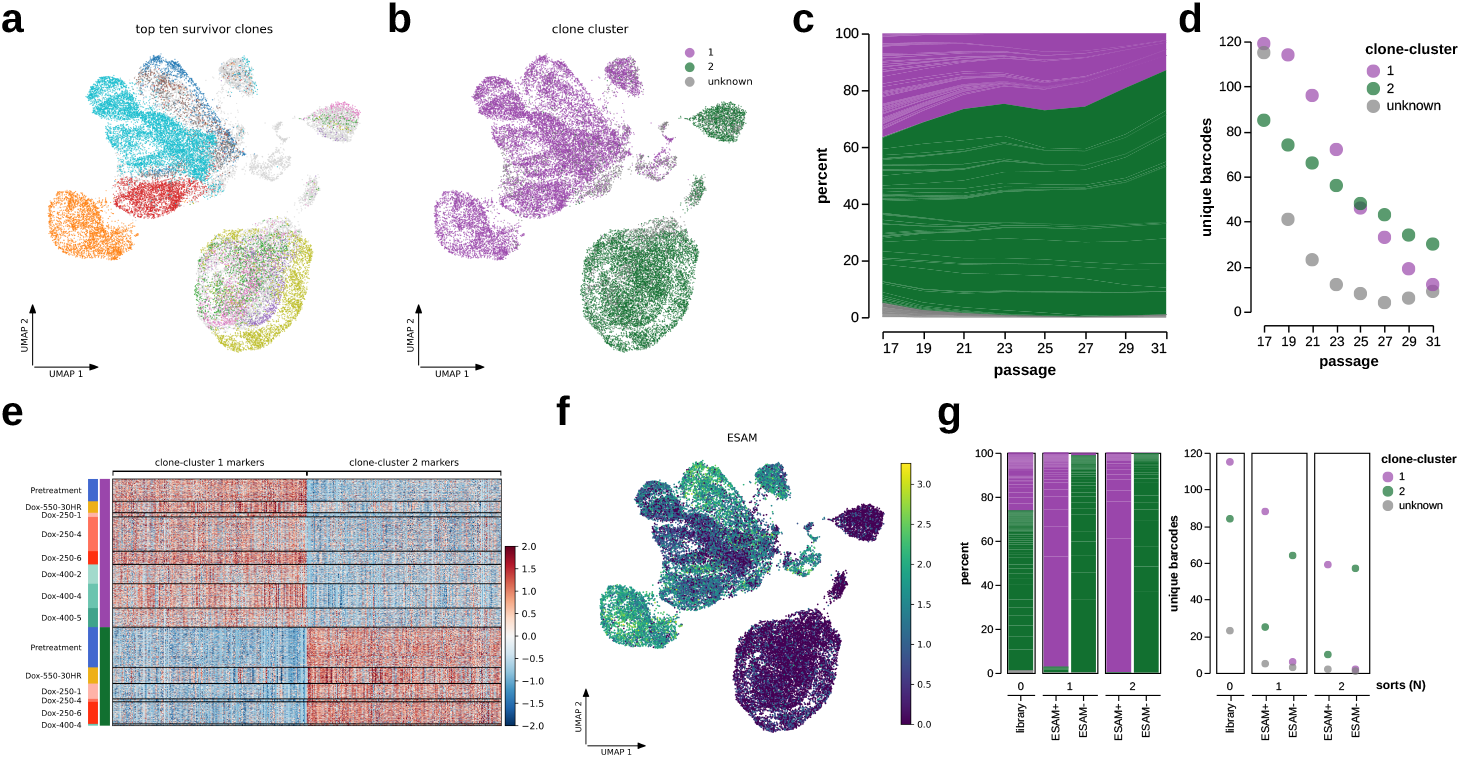
Barcoded MDA-MB-231s are comprised of clonally distinct subpopulations. **a)** UMAP of scRNA-seq annotated by the top ten survivor clones. **b)** UMAP of scRNA-seq annotated transcriptomically distinct clonal subpopulations 1 (purple) and 2 (green). **c-d)** Composition of total population by clone-cluster identity over time. **c)** Percent of clone-cluster identity within cell library over time. **d)** Total unique clones within cell library over time. **e)** Scaled expression of marker genes distinguishing clone-cluster 1 and 2, samples are separated into clone-cluster specific populations. Expression is scaled around 0 for each gene. **f)** UMAP of scRNA-seq annotated by normalized expression of ESAM. **g)** Composition of clone-cluster identity within cell library and serially sorted populations by ESAM expression. Percent of clone-cluster identity within cell library and sorted populations following successive cell sorting (left). Total unique clones within cell library and sorted populations (right).

### There is no differential expression within clone-clusters between high and low survivorship clones

Given the selection of only a subset of clones from doxorubicin treatment we then asked whether it was possible to identify a transcriptional signature that characterizes a pre-existing survivor population, consisting of cells primed to resist or recover from treatment. To distinguish clonal responses, we assigned a classification of “low” or “high” survivorship. Clones with an average percent change in total abundance of a clone across all treated samples >5% were assigned to the “high” survivorship category while clones with a percent change <5% (or that were not detected in any post-treatment replicates) were considered to have “low” survivorship potential. Of the surviving populations, there was no consistent bias toward either clone-cluster 1 or clone-cluster 2 (Fig. S1). This result is not surprising given the similar sensitivity of each subpopulation to doxorubicin when tested in isolation (Fig. S5). We then analyzed gene expression of high vs. low survivorship clones within each clonecluster in Pretreatment and Dox-550-30HR. There were no differentially expressed genes (DEG) passing cutoffs of (abs(log_2_FC) > 0.2 and FDR < 0.1) (Fig. S5a-d). Qualitative comparison of the top 5 differentially expressed genes showed little difference between populations at either time point. (Fig. 3a-d). Within each comparison there were a small number of DEGs with low p-values (<0.05) but they failed to pass the FDR cutoff criteria for significance (Fig. S5a-d). This observation suggests that clones persisting post-treatment do not possess an inherent survival advantage over those eliminated. To rigorously test this hypothesis, we conducted a permutation analysis by randomly shuffling the high and low survivorship classifiers and again found no significant differential expression (Fig. S6a-h). Although no significant differential expression was found between high and low survivor clones, substantial variability in final recovered clonal abundance was nonetheless observed.

**Fig. 3.**
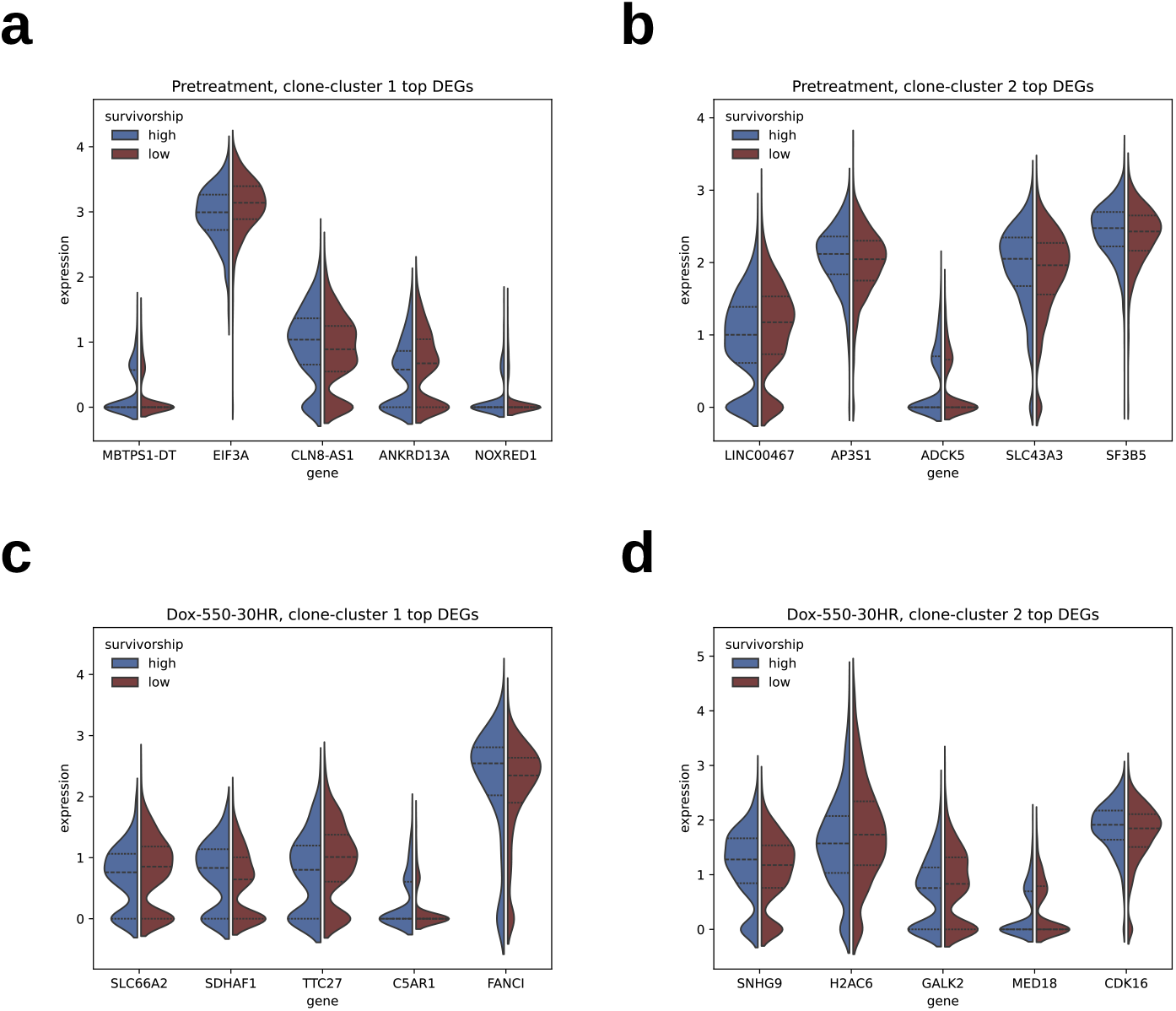
Clones that persist through selection do not display a pre-existing resistance signature compared to non-survivors. Normalized expression of top 5 differentially expressed genes between low and high survivorship within each clone cluster in Pretreatment (**a)** 1; **b)** 2) and Dox-550-30HR (**c)** 1; **d)** 2)

### Barcoded MDA-MB-231 subpopulations share an acute response to doxorubicin

Given the existence of distinct clone-cluster subpopulations, we then asked whether their acute response to doxorubicin was similar or distinct (Fig. 4a). We computed differential gene expression of each clone-cluster individually between Pretreatment and a high-dose on-going treatment, Dox-550-30HR (Fig. S7a,b). We compared the log_2_(fold change) of DEGs for each clone-cluster and found a positive correlation (R^2^ = 0.64, Fig. 4b). This consistent change in top marker genes between Pretreatment and Dox-550-30HR did not persist following recovery. The expression of top up- and down-regulated marker genes across all samples shows clone-cluster populations undergoing changes during treatment but then returning to a transcriptomic state more comparable to Pretreatment (Fig. 4d). Gene-set enrichment analysis with GO Biological Process terms on each individual set of DEGs showed an increase in DNA processing pathways (Mismatch Repair (GO:0006298), DNA Metabolic Process (GO:0006259)), and a decrease in pathways related to the electron transport chain (NADH Dehydrogenase Complex Assembly (GO:0010257)) and protein assembly (tRNA Aminoacylation for Protein Translation (GO:0010257), Cytoplasmic Translation (GO:0002181)) (Fig 4c). Thus despite the gene expression differences that distinguish that clone-cluster 1 and 2, there is significant overlap in their acute transcriptomic responses to doxorubicin.

**Fig. 4.**
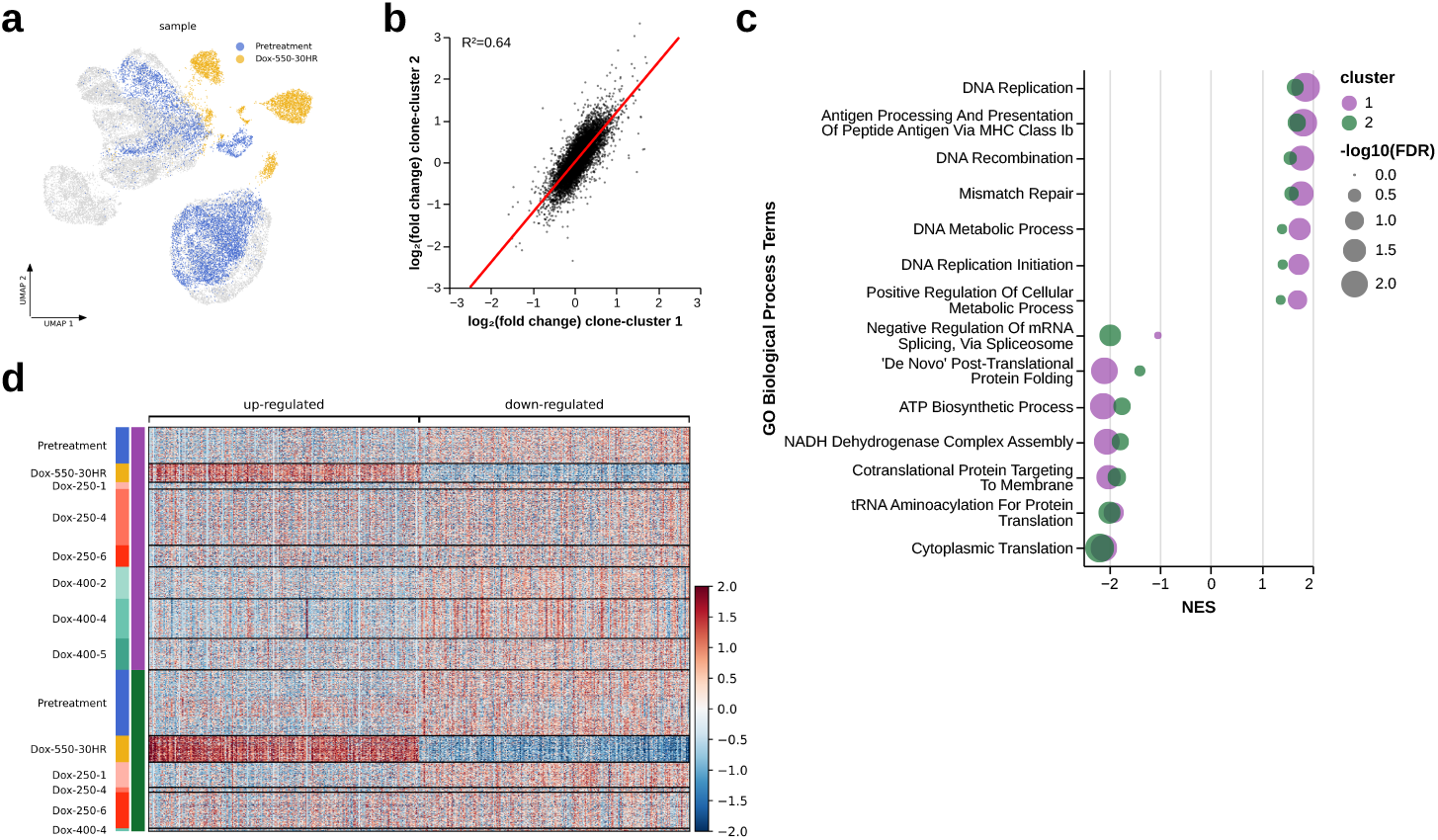
Acute response to doxorubicin has shared transcriptomic signature across subpopulations. **a)** UMAP of scRNA-seq colored by two samples Pretreatment (blue) and Dox-550-30HR (orange) **b)** Log_2_(Fold Change) of all differentially expressed genes between Pretreatment and Dox-550-30HR for each respective clone-cluster. **c)** Gene set enrichment analysis of GO Biological Process terms shared between both clonal clusters between Dox-550-30HR and Pretreatment. **d)** Scaled expression of marker genes (n=500) between Dox-550-30HR and Pretreatment across samples and clone-cluster subpopulations Expression is scaled around 0 for each gene.

### Individual clones can exhibit separate cell-states following recovery from acute doxorubicin treatment

Given the survival of some clones in more than one sample we asked how response to treatment affected their transcriptomic identity. To do this we assessed the differentially expressed genes of three individual clones (bg010 in Dox-250-4 and Dox-400-4 (Fig. 5a; Fig. S8a), bg032 in Dox-250-1 and Dox-250-6 (Fig. 5d; Fig. S8b), bg016 in Dox-250-4 and Dox-400-5 (Fig. 5g, Fig. S8c)) between Pretreatment and their respective post-treatment samples. Each clone was observed in more than one post-treatment sample (greater than 15 cells each); two clones (bg010, bg016) were found in samples of different drug doses (Fig. 5a,g). For all clones, we identified up and down-regulated genes (log_2_(fold change) > 0.2 and FDR < 0.05) shared between both of its samples and found small overlap compared to the total number of DEGs within each population (Fig. 5b,e,h). We additionally compared the enrichment of GO Biological Process terms within these post-treatment sample clones (5c,f,i) and found either no overlap in positive/negatively-enriched terms or, in the case of one clone (bg010), we found both positive enrichment and negative enrichment within the same terms (5c) depending on the recovered sample.

**Fig. 5.**
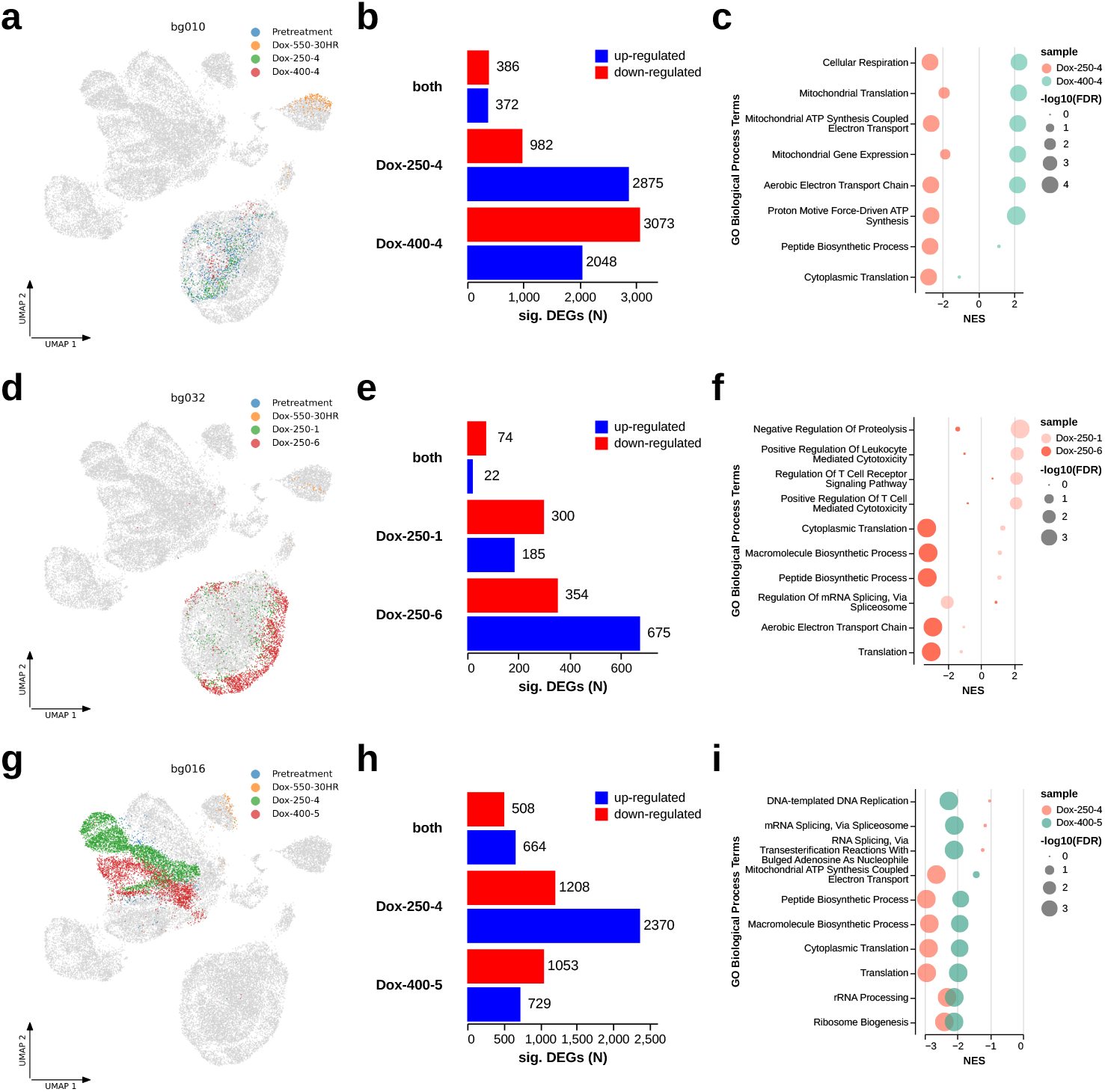
Clones undergo unique response to selection and subsequent recovery. Divergence of transcriptomic state was assessed for three clones: bg010, bg032, and bg016. **a**,**d**,**g)** UMAP of all cells highlighted by individual sample for respective clone. **b**,**e**,**h)** Total significantly up and down-regulated genes for each sample and shared between post-treatment samples (“both”) within clones (log2(FC) > .2, FDR < 0.05). **c**,**f**,**i)** Top enriched GO Biological Process terms by FDR for respective clones in Pretreatment vs post-treatment samples.

### Clonal populations that undergo high selection pressure have unique signatures

We then sought to determine the extent to which selection pressure induced *de novo* cell states within the population. Differential gene expression analysis comparing post-treatment clones to their pre-treatment parental clusters revealed that 6 out of 13 recovered clone populations exhibited significant upregulation of genes previously undetectable in their parental antecedents (6b). A clonal population within Dox-400-5 had 20 up-regulated genes that were lowly detected within its parental clone-cluster population (<25%) (Fig. 6a). Many of the genes detected only in post-treatment samples were non-coding RNA (Fig. 6b). In Dox-400-4-bg149, we found markers of increased proliferation, PCSK6 (Fig. 6d) and LDLRAD4 and higher expression of known TNBC tumor suppressors, SAMD5 (Fig. 6a). Within Dox-400-2-bg053 and to a lesser extent Dox-400-4-bg0149 there is an increased expression of SIRLNT (Fig. 6f). Together we find that clinically relevant markers arise in clonal populations that survive doxorubicin, suggesting the emergence of novel and heterogeneous cell states following treatment.

**Fig. 6.**
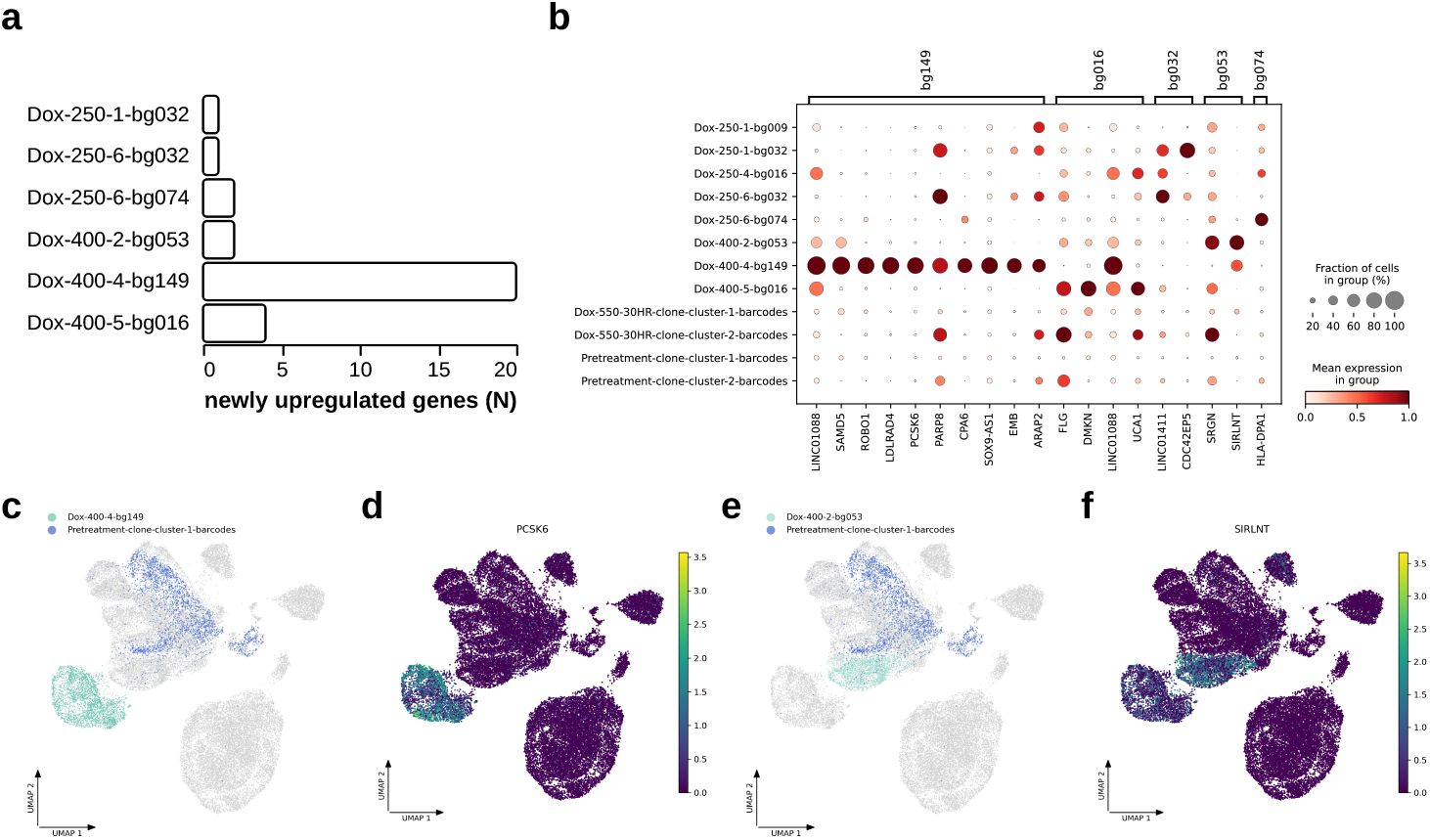
High doses of doxorubicin reveal clones specific transcriptomic signatures previously not detected. **a)** Number of significantly up-regulated DEGs detected in >60% of post-treatment sample cells and <25% of Pretreatment clone-cluster cells. **b)** Top newly up-regulated marker genes in clonal post-treatment populations. Post-treatment samples are grouped by sample and clone while Pretreatment and Dox-550-30HR are grouped by clone-cluster. **c)** UMAP of Dox-400-4-bg149 and Pretreatment-clone-cluster-1-barcodes. **d)** UMAP of PCSK6 expression. **e)** UMAP of Dox-400-2-bg053 and Pretreatment-clone-cluster-1-barcodes **f)** UMAP of SIRLNT expression.

## Discussion

TNBC remains one of the most challenging breast cancer subtypes to treat. This is exacerbated by its aggressive nature and significantly higher intratumoral heterogeneity, leaving late-stage patients with few options. Given this, we sought to better understand how a model of TNBC responds to high doses of doxorubicin at clonal resolution.

Genetic barcoding with heritable tags is an invaluable tool for tracking underlying clonal populations over time and through perturbation. Importantly, these tags are independent of classical phenotypic measurements that would be unstable for longterm identification in evolving populations and are increasingly used in the context of cancer [22]. With ClonMapper tracking of individual clonal populations, we were able to directly measure changes in clonal abundance in TNBC following exposure to doxorubicin. Unsurprisingly, there is a steep drop in population diversity consistent with a high dose of treatment. We suspect that the skew in the clonal diversity of individual samples is a characteristic of the outgrowth after treatment, rather than a reflection of differential intrinsic resistance of individual clones. Indeed, we observe some surviving clones at both high and low abundance among post-treatment replicate samples, which indicates that population dynamics or the presence of faster growing clones may dictate which surviving clone becomes the most abundant. The dynamics of treatment recovery were additionally confounded by *in vitro* culture conditions, such as passaging, which may have hampered the early recovery of senescent clones. Despite the commonality of some clones across samples we did not detect a common transcriptomic signature prior to treatment that would suggest specific populations were distinct in their survival. In limited approximations of individual barcode rate we found that high dose populations were more likely to harbor high-growth clones. Follow-up work should develop a more robust mathematical model of outgrowth of individual clones, accounting for differences in the intervals between treatment and initial recovery and considering the confounding effects of a local initial expansion.

Here and elsewhere [21, 23–25] there is evidence of two distinct clonal subpopulations within the MDA-MB-231 cell line. We found persistent survivor clones arising from both of these subpopulations, suggesting that they do not differ in resistance to doxorubicin. Follow-up studies of isolated subpopulations additionally supported the notion that they have similar potential to survive treatment. ScRNA-seq and subsequent surface-marker staining FACS revealed that these populations are clone-specific and they remain distinct transcriptomically during and following recovery from drug treatment. Its unclear the extent to which these subpopulations represent subtypes within TNBC or if they are unique to the MDA-MB-231 cell line.

Despite the lack of any pre-existing resistance in clones, we have identified sample- and clone-specific responses of TNBC to doxorubicin. Through mapping of transcriptomic state to individual clones we were able to observe unique, and at times divergent, phenotypes of clones as a result of treatment. Better understanding this stochasticity and response to treatment at a clonal resolution will be crucial in addressing the complex and unique clonal compositions of patient tumors. ClonMapper enables us to rapidly and iteratively perturb clonal populations to understand these responses. Designing new treatment strategies will require enhanced knowledge of the mechanisms governing this clonal diversification and the extent to which a clone can be steered [26–28].

Another key finding is the presence of previously undetected gene expression states in high-dose treatment samples. One sample in particular, Dox-400-4, up-regulates many genes which were not detected in other clones before or after treatment. One of these, LDLRAD4, is a negative regulator of TGF-beta signaling, a pathway associated with worse prognosis in gastrointestinal tumors [29] and increased tumorigenicity in hepatic cancers [30]. Another gene identified in Dox-400-4, PCSK6, in both knock-down [31] and exogenous treatment [32] has been shown to increase proliferation and invasiveness of MDA-MB-231 cells. This could explain why clone bg149 outcompeted other survivors within the Dox-400-4 sample. Separately, in two clones, we find consistent expression of lncRNA’s across post-treatment cells that were lowly expressed or undetected in pre-treatment populations. Many of the detected lncRNA in clone bg149 of Dox-400-4 are under characterized and their significance is hard to determine in this context. One lncRNA, SIRLNT, expressed in both bg149 and bg053, shas been associated with poor prognosis in breast cancer patients [33]. It is possible that some of the change in lncRNA expression is due to genetic alterations occurring during DNA repair in response to treatment. Follow-up studies would benefit from paired scDNA-seq to better characterize genetic changes within surviving clones.

The scope of this study is limited by its application within a sub-sampled population of a cell line model of TNBC and previous efforts have raised concerns about cell line variability [34] and clinical relevance [35]. It is not yet clear how these clone and subpopulation-specific behaviors would translate to more clinically relevant patient-specific models of TNBC, however it is possible that the rare cell state changes observed within these clones may be analogous to rare populations that emerge in patient tumors. It is also possible that the clonal survivorship observed in this model may depend heavily on specific *in vitro* conditions, not typical of the in vivo tumor microenvironment. Additionally, not well reflected here is the wealth of interactions that can occur *in vivo* through contact with immune and stromal populations. Incorporating these additional parameters may bolster our ability to predict how individual clones are likely to survive or evolve following treatment. Future work should focus on applying clonally resolved observations to patient-derived models of TNBC, which more closely resemble their starting tumor populations. This work serves as a basis for studying clonal behavior within TNBC as it relates to other drug treatments, as well as tumor progression and metastasis. Finally, this work highlights the importance of understanding clonal heterogeneity and the need for personalized strategies for unique patient tumors.

## Methods

### Cell culture

MDA-MB-231 cells were cultured in high glucose DMEM (Sigma D5796) supplemented with 10% FBS (Sigma, F0926-500mL) and 1X penicillin-streptomycin (Ther-moFisher, 15140122). Cells were passaged with 0.05% Trypsin-EDTA (ThermoFisher, 25300062) for 5 minutes.

### Generation of ClonMapper barcode plasmid library

Plasmid was prepared as previously described [18, 36]. Briefly, two oligonucleotides were ordered from IDT (CROPseq-PrimeF-BgL-BsmBI: GAGCCTCGTCTCC-CACCGNNNNNNNNNNNNNNNNNNNNGTTTTGAGACGCATGCTGCA and CROPseq-RevExt-BgL-BsmBI: TGCAGCATGCGTCTCAAAAC) and used in a 4X extension reaction to generate the barcode library insert. This insert was then ligated into a pre-digested backbone, Crop-Seq-BFP-WPRE-TS-hU6-BsmBI (Addgene, 137993), in a 50X golden gate reaction with BsmBI (NEB, R0739S). Following golden gate, plasmid was electroporated into *E. coli* in 500ml of 2xYT medium containing 100 ug*/*ml carbenicillin and incubated overnight at 37^*°*^C. Bacterial cells were pelleted by centrifugation at 6,000 RCF at 4^*°*^C for 15m and plasmid DNA was extracted using a QIAGEN Plasmid Plus Midi kit (QIAGEN, 12943).

### Generation of clonMapper barcoded MDA-MB-231 cells

Lentivirus was prepared as previously described [18, 36]. Cells were transduced with a low multiplicity of infection (MOI) of 0.1 to limit multiple integration events. 48h following transduction, BFP+ cells were plate sorted into a 96 well at varying concentrations 1 × 10^3^-1 × 10^4^ cells per well. Individual barcoded MDA-MB-231 cell libraries were expanded to approximately 5 × 10^6^ cells and sampled to confirm uniqueness and diversity skew of population. A barcoded MDA-MB-231 cell library (Pretreatment) from 1 × 10^3^ sorted cells was selected for further experiments.

### Drug treatment

Barcoded MDA-MB-231 cells were split into 24 replicate populations seeded at 5×10^4^ cells per well in a 24 well plate. After 24h, medium was replaced with cell-culture medium containing increasing concentrations of doxorubicin, 250, 400, 550 nM. Following 48h of exposure to drug, medium was replaced with fresh medium. Individual populations were then maintained separately and serially passaged into larger vessels and maintained in culture into they reached approximately 5 × 10^6^ total cells. Cells were pelleted for subsequent barcode abundance sampling and remaining cells were archived.

### Targeted barcode amplification

Genomic DNA barcodes were amplified and sequenced as previously described [18, 36]. Briefly, genomic DNA (gDNA) was harvested from ~1 × 10^6^ cells using the Invitrogen PureLink Genomic DNA kit (ThermoFisher, K182001). Barcodes were amplified from gDNA using a 2-step reaction. In the first reaction 2 ug of gDNA was amplified with primers flanking the barcode. A second PCR reaction was then carried out with 4 ng of the purified stage 1 reaction with primers containing Illumina adapters and indicies. All PCR reactions were cleaned with a right-sided cleanup using a 0.7-1.6x ratio of Ampure XP Beads (Beckman Coulter, A6388). Indexed samples were sequenced with either a NovaSeq 6000 (S1 flow cell, single read, 100 cycles) or NextSeq 500 (paired end, 75 cycles) by the UT Genome Analysis and Sequencing Facility.

### Barcode sequencing analysis

Barcode sequences were extracted and corrected from targeted amplification sequencing using `pycashier` [37]. Paired-end sequencing reads were first merged into consensus sequencing using `pycashier merge` with default settings. Next, barcode sequences were extracted from merged paired-end and single-read sequences using `pycashier extract` with default settings. Barcodes were ranked by initial abundance measurements from the cell library and were given an associated identifier in the format bg###. All other downstream targeted barcode abundance data was filtered based on presence within the earliest measured population. Pretreatment and all posttreatment samples were normalized to total BFP expression. One sample, Dox-400-5, was ~100% BFP resulting in no detected barcodes.

### Approximating clonal growth rate

To approximate growth rates of individual clones we assumed a population doubling time of 27 hours and calculated total cell number for the time of 2 passages (5 days/-passage). We multiplied these total cell numbers by individual barcode percentages for all two passage growth windows. These abundances were then fit to an exponential growth *N* (*t*) = *N*_0_*e*^*rt*^ to compute growth rates for each individual clone that was detected in at least 2 passages.

### Single-cell RNA-seq library preparation

To prepare scRNA-seq library, cryopreserved samples were thawed and cultured as described above. All samples were passaged at lease once to allow recovery from cryopreservation. Libraries were prepared in two batches (batch 1: Pretreatment, Dox-550-30HR, Dox-250-4 and Dox-250-6 and batch 2: Pretreatment, Dox-250-1, Dox-400-2, Dox-400-4, Dox-400-5). To generate Dox-550-30HR, prior to sample harvest Pretreatment was split and treated with 550nM starting 30h before harvest. All samples were detached from plates and resuspended in PBS with 0.04% BSA (ThermoFisher, 15260037). Libraries were generated using a 10X Chromium and Single Cell 3’ Gene Expression (v3.1) Reagent Kit with a loading target in each well of the Chromium of 10,000 cells. All library preparation steps were carried out by by the UT Genome Analysis and Sequencing Facility according to the manufacturer’s instructions.

### Clonally-resolved scRNA-seq analysis

All single-cell data was aligned to GRCh38 (refdata-gex-GRCh38-2020-A, 10X Genomics) and processed using cellranger (v8.1). Raw filtered counts for each sample were first generated with cellranger count. Downstream analysis was carried out using python and scanpy [38] (see Code Availability). Briefly, all samples were merged into a single object containing raw counts and associated metadata, combining Pretreatment from each batch into a single sample. Samples were further processed according to single cell best practices from the Theis Lab [39]. Briefly, data was filtered by thresholding on total counts, number of genes, and percent counts in the 20 genes using a median absolute deviation of 5 for any category. Cells were additionally filtered by a percent mitochondrial gene count with a median absolute deviation less than 3 or a percent mitochondrial gene above 20%. Genes were filtered if not detected in greater than 20 cells. Doublets were identified and filtered using scDblFinder [40].

Counts were normalized using a log-shifted normalization as described by single cell best practices [39]. We performed dimensionality reduction by finding highly variable genes, calculating principal components, computing nearest neighbors and then finally a UMAP representation [41]. To generate clonal barcode assignments, we first generated sam files containing only unmapped reads from the sam file generated for each sample by `cellranger count`. We extracted individual sequences using `pycashier` scrna with a minimum length of 10bp and a maximum length of 21bp, then matched these to to the start and end of any sequence that was 19-21bp. We then removed any remaining sequences less the 19bp and found in cells that did not pass the aforementioned expression-based filtering. Sequencing and amplification errors were corrected by performing message-passing clustering with starcode [19]. These corrected barcodes were than paired to individual cells. Clone-clusters were computed through hierarchical clustering with a cutoff threshold of 225.

### Differential expression and pathway analysis

Unless otherwise stated differential expression analysis was limited to cells assigned a single barcode. Differentially expressed genes were computed using MAST [42]. Comparisons including more than one clone incorporated clonal identity as a random effect using a mixed-effect model, whereas within clone comparisons used a generalized linear model. We then adjusted for multiple comparisons using a Benjamini-Hochberg correction. Unless otherwise noted genes were considered differentially expressed if log_2_(fold change) > 0.2 and FDR < 0.05. To determine enrichment of gene expression pathways we used Preranked GSEA with gseapy [43]. Genes were ranked by taking the sign of their log_2_(fold change) and multiplying it by −log_10_(p-val)

### Subpopulation separation by ESAM expression with FACS

Barcoded MDA-MB-231 cells were stained with anti-ESAM-FITC (Miltenyi Biotec, 130-130-697) and FACS sorted into two populations ESAM+/BFP+ and ESAM-/BFP+. This procedure was repeated with previously sorted populations for further purification of expected populations, i.e. ESAM+/BFP+ cells were purified from gated ESAM+/BFP+ expression and ESAM-/BFP+ cells were again gated from ESAM-/BFP+ cells. From each of the following timepoints, prior to sort, after first sort, and after second sort, gDNA was harvested from pelleted cells. Unique barcode abundance was confirmed in each population as described above.

## Data Availability

Targeted barcode sequencing data has been deposited in the Gene Expression Omnibus (GEO) under accession code GSE291679. Single cell RNA-sequencing was deposited under accession code GSE291678

## Code Availability

Code for analysis and figure generation can be found on the Brock Lab GitHub at brocklab/clonmapper-tnbc-doxorubicin.

## Acknowledgements

We are grateful for funding from the NIH R01CA255536 and U01CA253540 (to A.B.) and and F31CA268833 (A.L.G). We thank the core facilities at UT Austin for access to their services and knowledge. Sequencing was performed by the Genomic Sequencing and Analysis Facility at UT Austin, Center for Biomedical Research Support (RRID: SCR_021713). Flow cytometry and FACS was performed at the Center for Biomedical Research Support Microscopy and Imaging Facility at UT Austin (RRID: SCR_021756).

## Author Contributions

D.M.: Conceptualization, Methodology, Software, Formal Analysis, Data Curation, Writing—Original Draft, Writing—Review & Editing, Visualization. A.L.G.: Methodology, Writing—Review & Editing. A.B.: Resources, Writing—Review & Editing, Supervision, Project administration, Funding acquisition.

