## Supplementary Figures for "Lineage Tracing Reveals Clone-Specific Responses to Doxorubicin in Triple-Negative Breast Cancer"

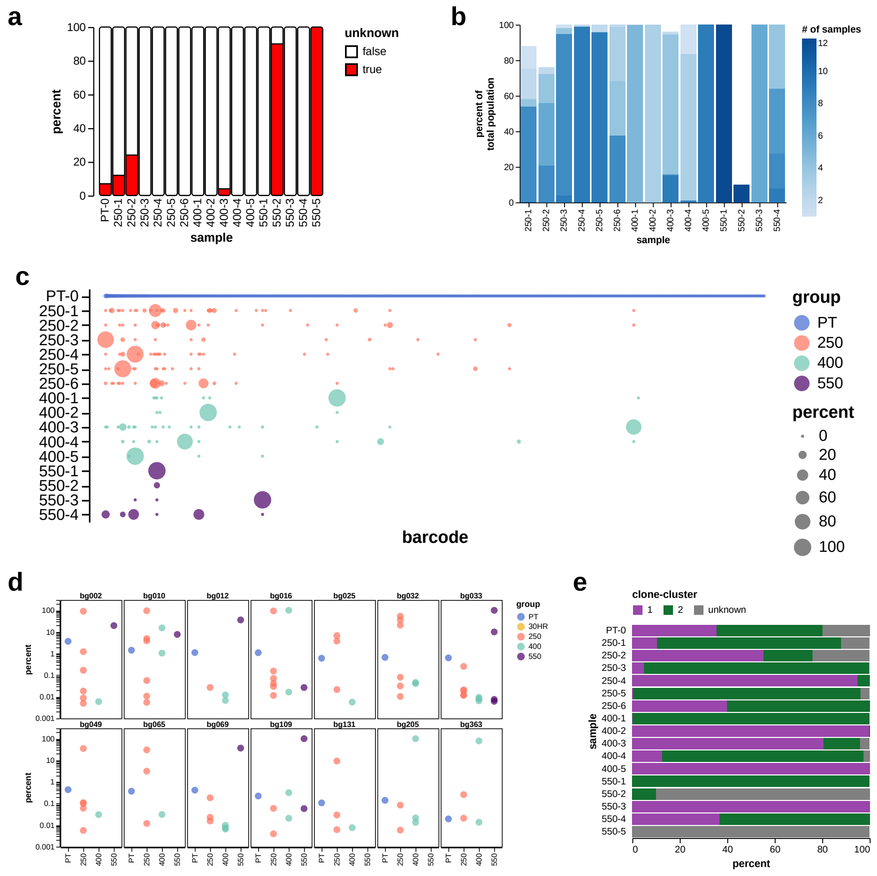


**Supplementary Figure S1: Generation of barcoded TNBC population. a)** Percent of barcoded populations with no clonal identity as determined by expression of BFP. **b)** Percent of each population colored by total of samples it's underlying clones share. **c)** Total abundance of all barcodes across cell library and post-treatment samples. Barcodes are ordered by initial abundance in PT. **d)** Top 15 barcodes by cumulative Abundance in post-treatment samples. Each panel is total percent of a single clone within each sample, colored by group. **e)** Percent total abundance in Pretreatment and post-treatment samples of barcodes by clone-cluster.

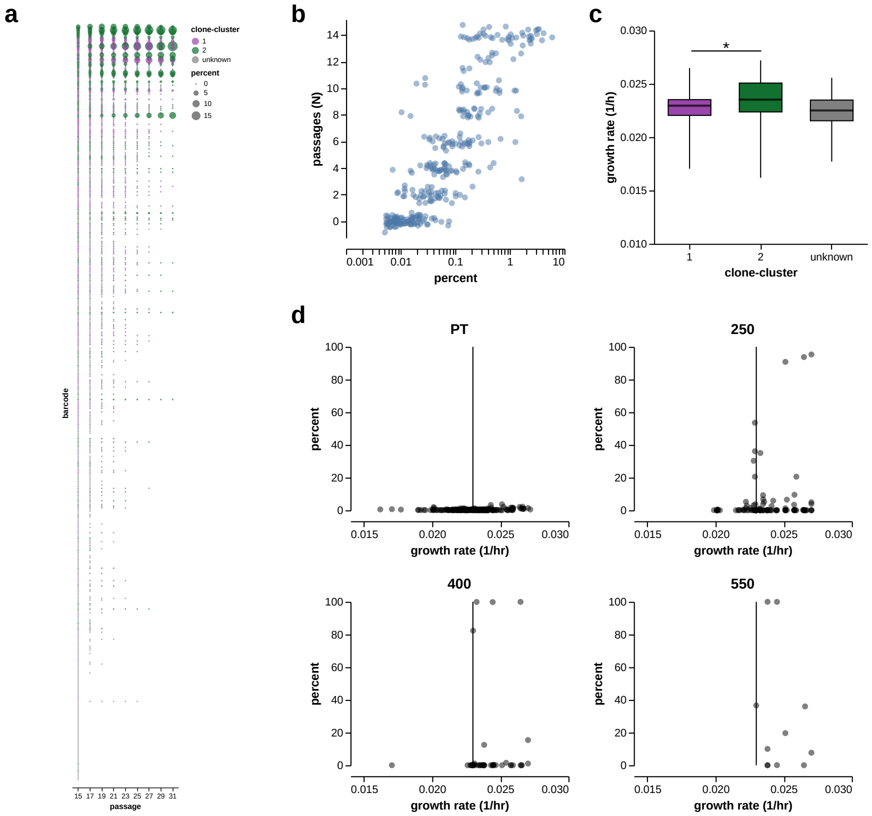


**Supplementary Figure S2: Clonal Growth in barcoded TNBC population.** **a)** Dotplot of total abundance across cell library maintained in culture. Barcode is ordered by abundance at passage 15. **b)** Number of passages from starting that a clone was detected in compared to it's starting abundance when cell library was maintained for 14 passages. **c)** Approximate growth rates for barcode by clone-cluster. P-value 0.00135; whiskers, min and max values; box, first and third quartile; line, median. **d)** Approximate growth rates of barcodes and abundance in each sample by group (PT: Pretreatment, 250,400,550: Doxorubicin does in $\mathrm{nM}$). Line, median growth rate for all barcodes.

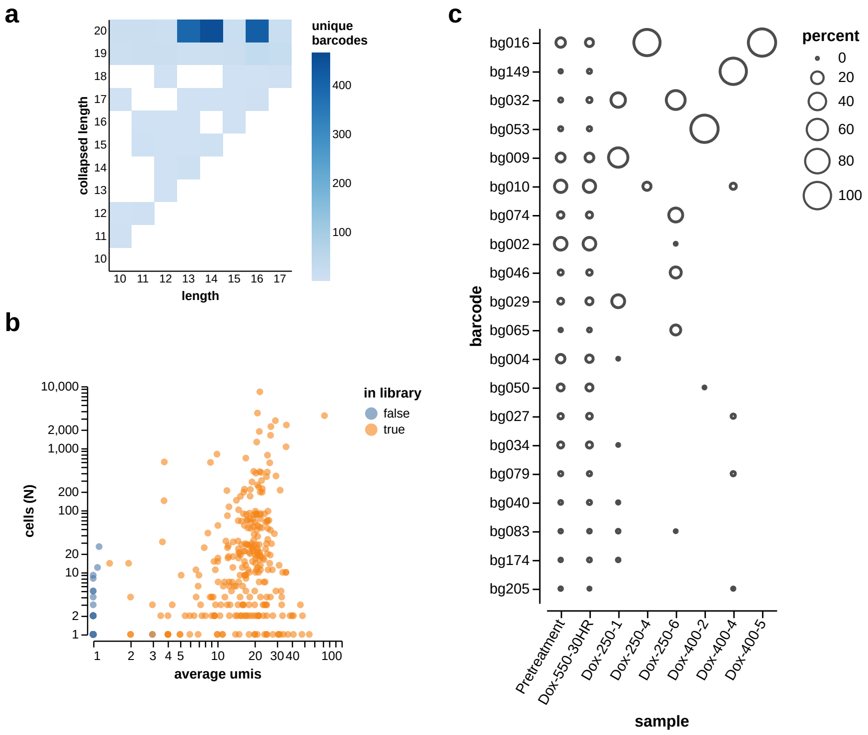


**Supplementary Figure S3: Clonal populations at single-cell resolution. a)** Heatmap of the number of barcodes collapsed into larger sequences. **b)** Barcode sequences from scRNA-seq plotted by average umis and total cell colored based on their detection in the targeted sequencing of initial cell library. **c)** Abundance of top post-treatment clones across all scRNA-seq samples. Barcodes are ordered by cumulative abundance across samples.

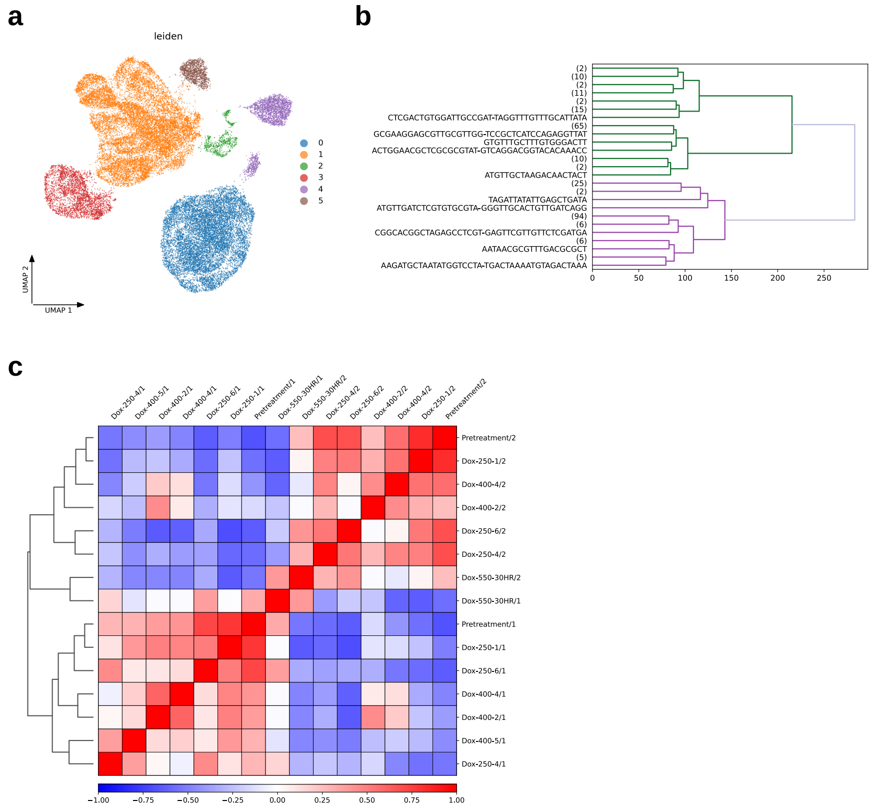


**Supplementary Figure S4: Clone Clusters in Barcoded MDA-MB-231s. a)** UMAP of scRNA-seq annotated by leiden cluster. **b)** Hierarchical clustering of Pretreatment scRNA-seq clones with split with threshold of 255. **c)** Correlation matrix of Pearson's Correlation Coefficient in samples split by clone-cluster subpopulations.

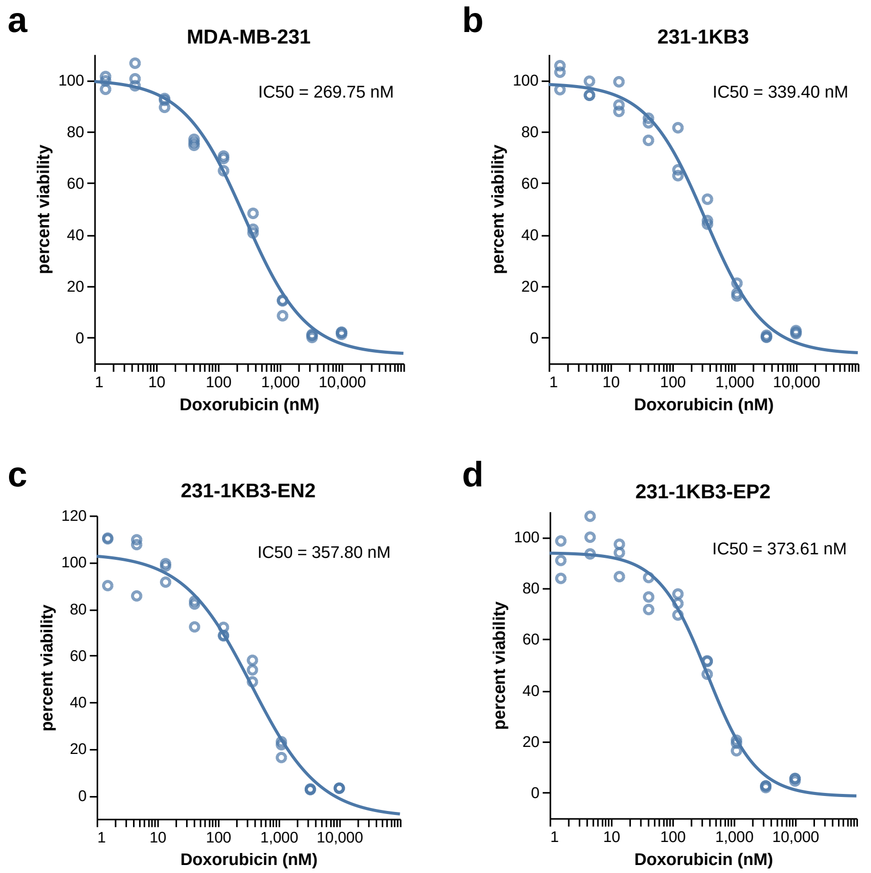


**Supplementary Figure S5: Response in MDA-MB-231 to doxorubicin treatment.** Dose responses curves of parental and barcoded MDA-MB-231 cell line: **a)** MDA-MB-231, **b)** 231-1KB3, barcoded MDA-MB-231 library, **c)** 231-1KB3-EN2, ESAM$^-$ barcoded MDA-MB-231, and **d)** 231-1KB3-EP2, ESAM$^+$ barcoded MDA-MB-231.

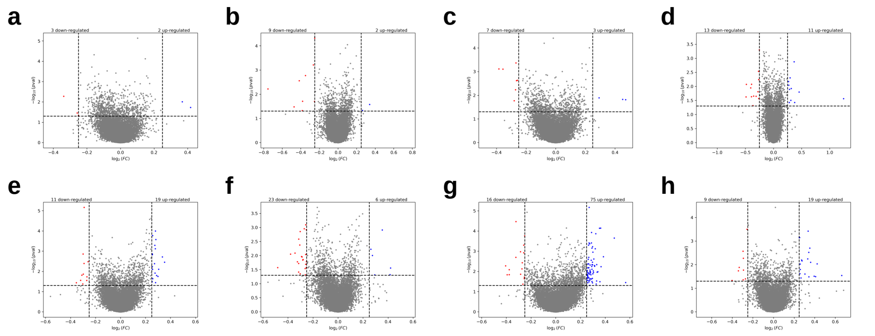


**Supplementary Figure S6: Clones that persist through selection do not display a pre-existing resistance signature compared to non-survivors.** Volcano plots of differentially expressed genes between low and high survivorship within each clone cluster in samples: Pretreatment (**a)** 1; **b)** 2) and Dox-550-30HR (**c)** 1; **d)** 2). Volcano plots of differentially expressed genes between low and high survivorship within each clone cluster in samples following random shuffling of survivorship classification: Pretreatment (**e)** 1; **f)** 2) and Dox-550-30HR (**g)** 1; **h)** 2).

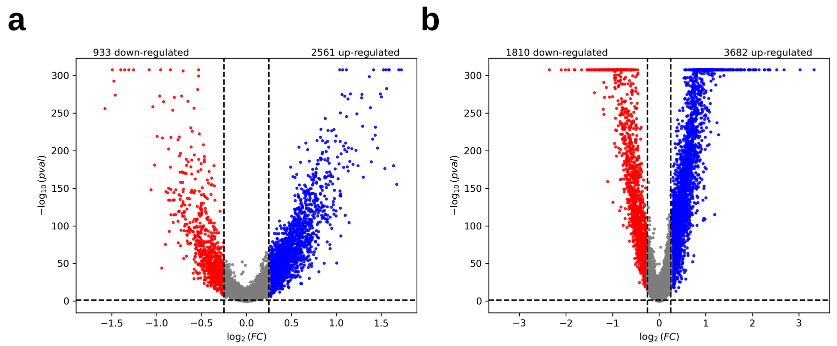


**Supplementary Figure S7: Doxorubicin induces differentially expressed genes across clonal subpopulations.** Volcano plots of differentially expressed genes between Pretreatment and Dox-550-30HR in **a)** clone-cluster 1 and **b)** clone-cluster 2.

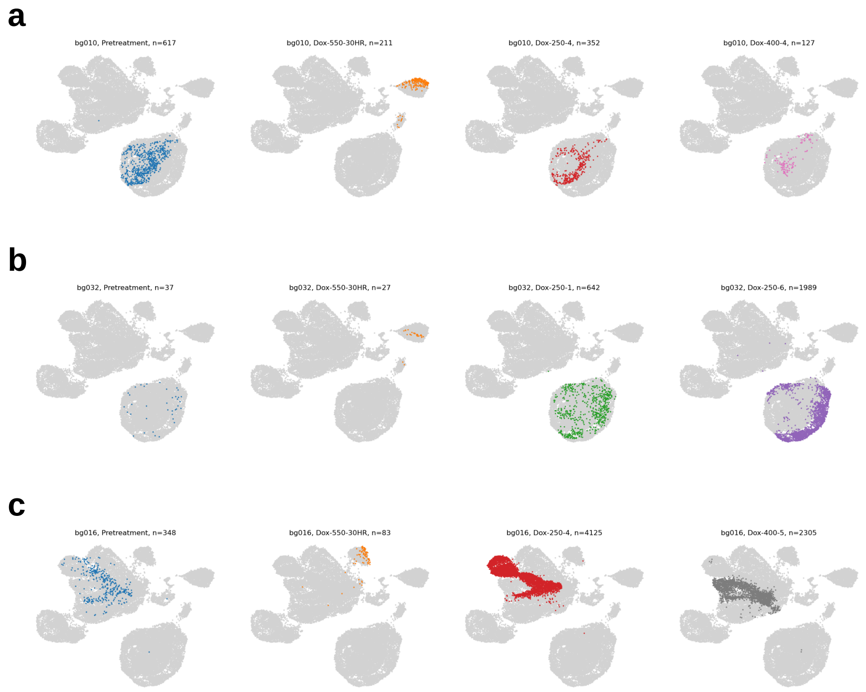


**Supplementary Figure S8: Divergent clonal populations following treatment.** UMAP of scRNA-seq separated by individual samples for **a)** bg010, **b)** bg032, and **c)** bg016.
